# Gene-modified NK Cells Expressing CD64 and Pre-loaded with HIV-specific BNAbs Target Autologous HIV-1 Infected CD4^+^ T Cells by ADCC

**DOI:** 10.1101/2024.06.20.599937

**Authors:** Costin Tomescu, Adiana Ochoa Ortiz, Lily D. Lu, Hong Kong, James L. Riley, Luis J. Montaner

**Affiliations:** The Wistar Institute, HIV Immunopathogenesis Laboratory, BEAT-HIV Delaney Collaboratory, Philadelphia, PA 19104; The Wistar Institute, Molecular Screening Facility, Philadelphia, PA 19104; The University of Pennsylvania, Perelman School of Medicine, Department of Microbiology, Center for Cellular Immunotherapies, BEAT-HIV Delaney Collaboratory, Philadelphia, PA, 19104

**Keywords:** High Affinity Fc Receptor CD64, NK Cells, CD57 Maturation Marker, Persons Living with HIV (PLWH), Antibody Dependent Cellular Cytotoxicity (ADCC), Broadly Neutralizing Antibodies (BNAbs)

## Abstract

NK cells can efficiently mediate antibody-dependent cellular cytotoxicity (ADCC) of antibody coated target cells via the low-affinity Fc-receptor, CD16, but cannot retain antibodies over time. To increase antibody retention and facilitate targeted ADCC, we genetically modified human NK cells with the high-affinity Fc receptor, CD64, so that we could pre-load them with HIV-specific BNAbs and enhance their capacity to target HIV infected cells via ADCC. Purified NK cells from the peripheral blood of Control Donors or Persons Living with HIV (PLWH) were activated with IL-2/IL-15/IL-21 cytokines and transduced with a lentivirus encoding CD64. High levels of CD64 surface expression were maintained for multiple weeks on NK cells and CD64 transduced NK cells were similar to control NK cells with strong expression of CD56, CD16, NKG2A, NKp46, CD69, HLA-DR, CD38, and CD57. CD64 transduced NK cells exhibited significantly greater capacity to bind HIV-specific BNAbs in short-term antibody binding assay as well as retain the BNAbs over time (1 week antibody retention assay) compared to Control NK cells only expressing CD16. BNAb pre-loaded CD64 transduced NK cells showed a significantly enhanced capacity to mediate ADCC against autologous HIV-1 infected CD4^+^ primary T cells in both a short term 3 hour degranulation assay as well as a 24 hour HIV p24 HIV Elimination Assay when compared to control NK cells. A chimeric CD64 enhanced NK cell strategy (<u>NK E</u>nhancement <u>S</u>trategy, “NuKES”) retaining bound HIV-specific antibody and targeted ADCC represents a novel autologous primary NK cell immuno-therapy strategy against HIV.

## Introduction

One of the foremost goals of HIV cure-directed gene therapy strategies is to eliminate virally infected cells with the greatest potential to produce replication competent virions, of which HIV infected cells expressing envelope are a critical subset [1–5]. In the peripheral blood, cytotoxic NK cells (CD56^dim^/CD16^+^) represent a critical innate immune effector cell type with the potential to eliminate envelope expressing cells coated with HIV-specific antibodies through the process of Antibody Dependent Cellular Cytotoxicity (ADCC) [6–10]. However, obstacles remain that prevent NK clearance of infected targets though ADCC when NK cells only utilize the traditional Fc Receptor, CD16. Firstly, due to the low-affinity nature of CD16 [11–15], cytotoxic NK cells are not capable of retaining HIV-specific antibodies long enough to kill virally infected target when they encounter the antibody before the target cell. Secondly, CD16 possesses a proteolytic cleavage site in the extra-cellular domain that allows it to be negatively regulated by the Matrix Metallo-proteinase ADAM-17, which cleaves CD16 off the surface of NK cells after degranulation and prevents sequential ADCC activity over time [16–18].

In the cancer Immuno-therapy field, several strategies have been developed to overcome these obstacles by genetically modifying NK cells with altered Fc receptors. This included efforts to remove the proteolytic cleavage site for ADAM-17 from the low-affinity CD16 Fc Receptor [19, 20], express the high-affinity Fc receptor, CD64, in tandem with CD16 [21], or engineer chimeric Fc receptors containing the extra-cellular portion of CD64 merged with the intra-cellular portion of CD16 [22]. As the high-affinity CD64 Fc receptor lacks the proteolytic cleavage site utilized by ADAM-17 to degrade CD16, NK cells expressing CD64 are resistant to the negative regulation of ADCC that traditionally plagues NK cells only expressing CD16 [22, 23]. Furthermore, due to the high affinity nature of CD64 for antibodies [24, 25], NK cells engineered to express CD64 (or chimeric CD64/CD16 Fc receptors) have been shown to possess the capacity to retain broadly neutralizing antibodies (BNAbs) specific for cancer ligands over long periods of time and mediate targeted ADCC against cancer cells both *in vitro* and *in vivo* [21, 23, 26]. However, these various Fc enhancement strategies were not explored on primary NK cells from the peripheral blood due to low lentiviral gene transduction efficiency typically associated with primary NK cells [27–31]. As a result, Fc enhancement strategies have been largely confined to NK cell lines or induced NK cells differentiated from pluripotent stem cells in heterologous situations where the NK cells and target cells are derived from different individuals [21, 23, 26].

*In vitro* NK functional assays using heterologous target cells (i.e.: when targets and NK effectors are from different individuals), may artificially inflate the target cell sensitivity to NK cell lysis due to the potential mismatch in MHC-I ligands and the resulting reduction in NK inhibitory receptor signaling regulatory signals that normally limit NK cell recognition of self [32–35]. By contrast, an autologous functional NK cytotoxicity assay retains physiologically relevant KIR/MHC-I interaction and better approximates *in vivo* NK effector cell activity against hard to kill targets [36–38]. Here, we investigated if we could adapt a CD64 Fc genetic enhancement strategy using lentivirus transduction of human NK cells isolated from the peripheral blood to target HIV-1. Our goal was to pre-load CD64 genetically modified NK cells with HIV-specific BNAbs and direct them to kill HIV-1 infected CD4^+^ Primary T cells expressing envelope in our previously described autologous assay system for measuring both NK-mediated direct cytotoxicity [37, 39] as well as ADCC [40, 41]. Here, we show that antibody pre-loaded chimeric CD64 NK cell strategy or “NuKES” (<u>NK</u> <u>E</u>nhancement <u>S</u>trategy) possess an enhanced capacity to retain HIV-specific BNAbs and mediate targeted ADCC against autologous CD4^+^ Primary T cells infected with HIV.

## Results

### High-level Lentiviral Transduction and Stable CD64 Expression on the Surface of Human NK cells Purified from the Peripheral Blood

With the goal of transducing NK cells isolated from peripheral blood with the full-length high-affinity Fc Receptor 1a, CD64, we overcame the low lentiviral gene transduction efficiency typically observed with primary cells [27–31] through strong pre-activation with a cytokine only (feeder-cell free) cocktail containing IL-2/IL-15/IL-21 and use of third generation lentiviral constructs [25]. As shown in **Figure 1A**, this approach led to strong CD64 surface expression on peripheral blood NK cells over multiple days that was not diluted out. The durable CD64 expression over time allowed for a long therapeutic window of several weeks to use these genetically modified NK cells in subsequent functional assays (**Figure 1B**). Importantly, we could reach high CD64 expression levels using both NK cells derived from small whole blood draws of control donors as well as larger Apheresis preparation from PLWH (**Figure 1C**).

**Figure 1.**
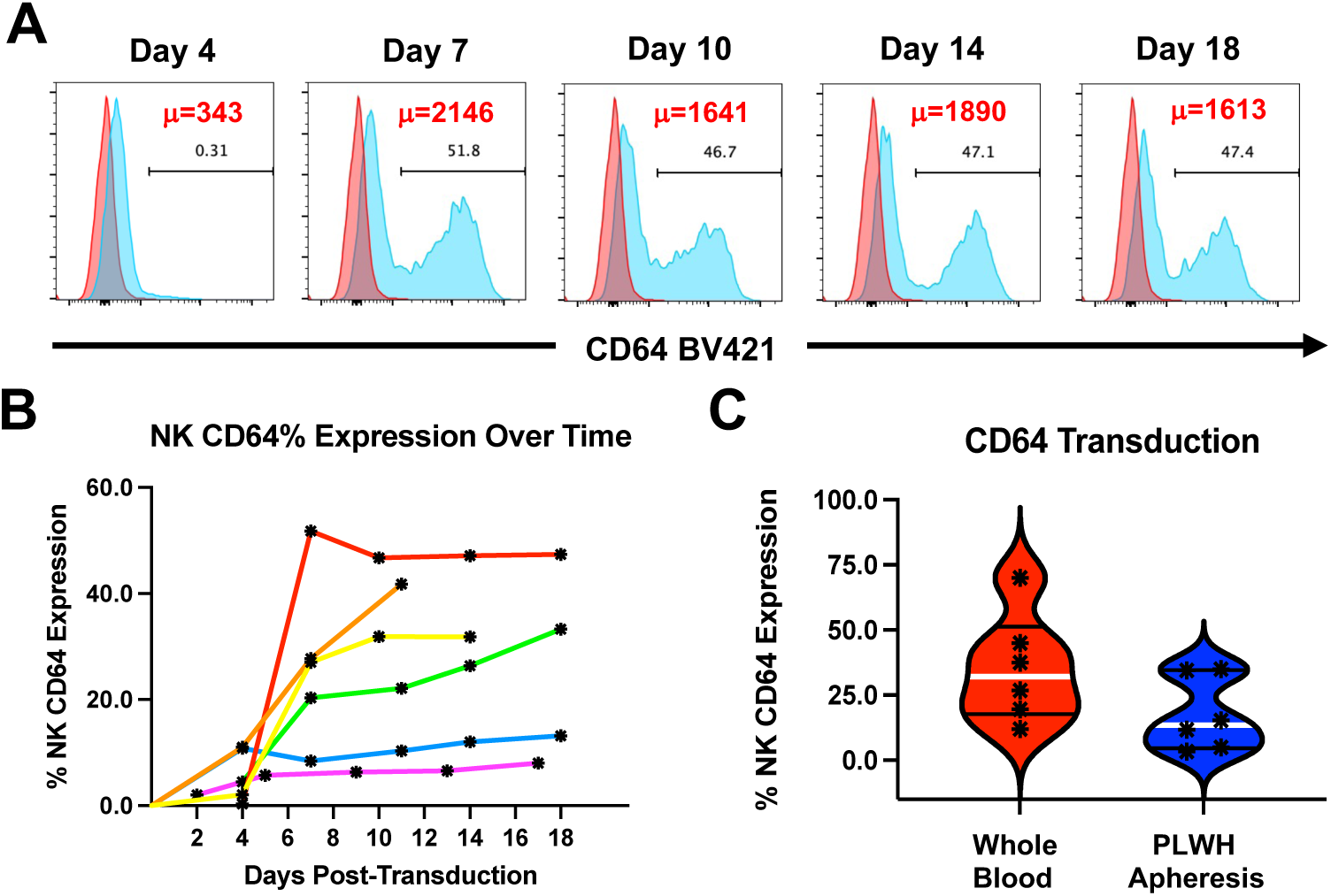
High-level Lentiviral Transduction and Stable CD64 Expression on the Surface of Human NK cells Purified From the Peripheral Blood. (A-B) NK cells were purified by negative magnetic bead separation (Miltenyi Corporation) to 99% purity from the peripheral blood of control donors or Persons Living With HIV, activated with 100 u/mL of IL-2, 100 ng/mL of IL-15 and 100 ng/mL of IL-21 for two days, and then either mock or virus transduced with CD64 expressing Lentiviral particles. NK cells were stained for Flow Cytometry every few days and CD64 surface expression was measured on CD56+/CD3-NK cells. (A) A representative experiment showing CD64 expression over time on Lentiviral Transduced NK cells (Blue Histogram) as compared to Control Non-Transduced NK cells (Red Histogram). The Geometric Mean Fluorescence Intensity of CD64 expression is shown in upper right-hand part of each histogram in red. (B) Composite graph showing CD64 expression over time on Lentiviral Transduced NK cells from six individual experiments. (C) Composite graph showing high CD64 transduction frequencies using both NK cells derived from small whole blood draws from control donors or large-scale Apheresis preps from PLWH.

Next, we determined the degree of CD64 transduction in mature CD57 NK cells. CD57 expression been associated with enhanced NK cytotoxic function [42–45] and NK cells expressing CD57 have been identified as the pre-dominant NK effector population in peripheral blood able to mediate higher ADCC against HIV-1 infected autologous CD4^+^ T cells in both control donors and PLWH [40]. As shown in **Figure 2A**, the CD57 positive NK cells from the peripheral blood were readily transducible with the CD64 lentivirus in our system. Importantly, we could maintain CD57 maturation status on our NK cells despite multiple weeks in culture (**Figure 2B**) and there was no difference (p=n.s., n=10) in the CD57 maturation status between CD64 transduced NK cells and control NK cells (**Figure 2C**). Of note, there was also no measurable difference in the expression of other NK phenotypic or activation markers including CD56, CD16, NKp46, NKG2A, CD38, and HLA-DR between CD64 positive and control NK cells (**Supplementary Figure 1**).

**Figure 2.**
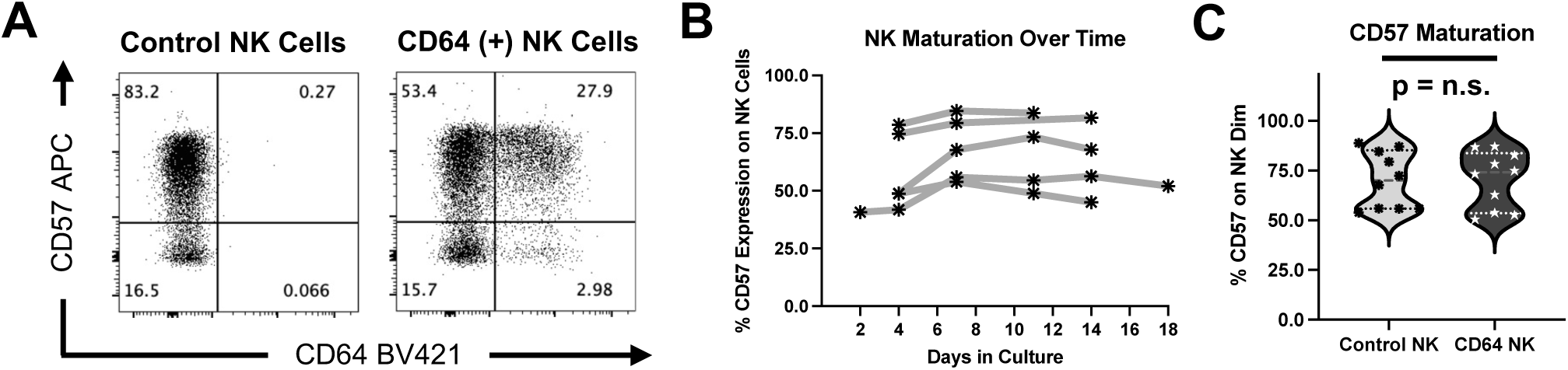
Successful Lentiviral Transduction of CD57 Mature NK Cells and Maintenance of Stable CD57 Maturation Status on Transduced NK cells Over Time. **(A-C)** NK cells were purified by negative magnetic bead separation (Miltenyi Corporation) to 99% purity from the peripheral blood, activated with 100 u/mL of IL-2, 100 ng/mL of IL-15 and 100 ng/mL of IL-21 and then either mock or virus transduced with CD64 expressing Lentiviral particles. NK cells were stained for Flow Cytometry every few days with fluorescently conjugated antibodies to NK Phenotypic Markers and CD57 surface expression was measured on CD56+/CD3-NK cells. **(A)** A representative experiment showing the successful CD64 Lentiviral Transduction of CD57 Positive Mature NK cells on Day 7 post transduction. **(B)**. Composite graph showing the stable expression of CD57 over time on Control NK cells cultured with our feeder cell free, cytokine only cocktail in five individual experiments. **(C)** Composite graph showing no significant difference in the CD57 Maturation Status of Control or CD64 Transduced NK cells at Day 7 post Transduction (n=10).

### CD64 Transduced NK cells Can Bind and Retain HIV-specific BNAbs To a Significantly Greater Degree Than Control NK cells

One of the purposes for genetically modifying NK cells with the high-affinity Fc Receptor CD64 was to pre-load them with HIV-specific BNAbs so they could seek out and destroy virally infected cells expressing envelope in an antigen specific fashion. To that end, we fluorescently conjugated the 10-1074 BNAb, specific for the V3 Glycan of HIV-1 envelope [46], with the FITC fluorochrome and pre-loaded it onto control and CD64 transduced NK cells in a short-term antibody binding assay. As shown in **Figure 3A and B**, CD64 transduced NK cells bound the FITC conjugated 10-1074 BNAb to a higher degree than control NK cells both in terms of frequency of initial NK cell binding and Geometric Mean Fluorescence Intensity (GMFI) of BNAb molecules being bound per NK cell. Importantly, every CD64 expressing NK cell (30% in the representative experiment shown) bound the FITC conjugated 10-1074 BNAb and became double positive whereas only a fraction of control NK cells could retain the bound BNAb after a washing step (**Figure 3A and B**). This difference in short-term HIV-specific BNAb binding between CD64 transduced NK cells and control NK cells was both significant (*p=0.0312*, n=6) and robust as the difference in Geometric Mean Fluorescence Intensity was on average over 10,000 units per NK cell (**Figure 3C and D**).

**Figure 3.**
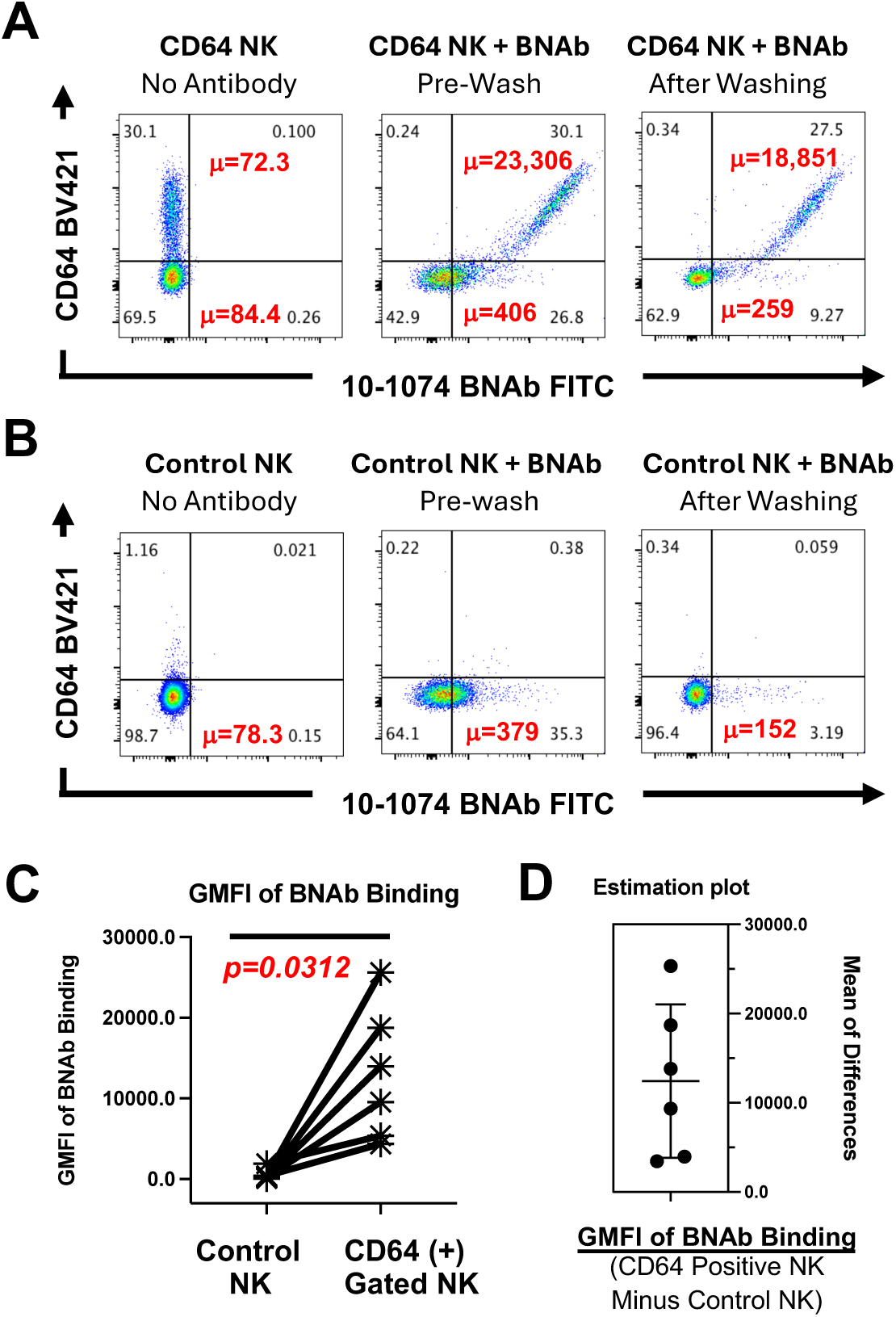
CD64 Transduced NK cells Can Bind HIV-specific BNAbs To a Significantly Greater Degree Than Control NK cells in a Short-term Assay of BNAb Binding. **(A-C)** Control and CD64 Transduced NK cells were prepared as in Figure 1. On Day 7 post transduction, Control or CD64 transduced NK cells were pre-loaded with 50 μg/ml of FITC Conjugated HIV-specific 10-1074 BNAb for 15 minutes. NK cells were stained for CD64 Surface Expression and HIV-Specific BNAb Binding by Flow Cytometry before and after washing to measure short-term BNAb retention. **(A-B)** A representative experiment showing the successful capture and short-term retention of HIV-specific BNAbs by **(A)** CD64 Lentiviral Transduced NK cells compared to **(B)** Control NK cells on Day 7 post transduction. The Geometric Mean Fluorescence Intensity of BNAb binding through CD64 and CD16 is shown in upper right-hand and lower-right hand part of each dotplot in red. **(C)** Composite graph showing statistically greater capacity of CD64 Positive Gated NK cells to retain HIV-specific BNAbs on the cell surface as reflected in Geometric Mean Fluorescence Intensity after washing in a short-term binding assay (n=6). **(D)** Estimation plot shows the difference in Geometric Mean Fluorescence of BNAb binding between CD64 Lentiviral Transduced NK cells and Control NK cells in the short-term binding assay after washing.

We next assessed the durability of BNAb binding by CD64 positive Transduced NK cells over time in a long-term assay of BNAb retention. Initially, we observed over 99% of transduced NK cells expressing CD64 bound the FITC conjugated 10-1074 BNAb and retained it overnight after washing (**Figure 4A**). By contrast, only a fraction of control NK cells expressing CD16 could bind the BNAb initially with a Geometric Mean Fluorescence Intensity that was substantially lower (GMFI=910 compared to GMFI=4,283) than the CD64 Transduced NK cells (**Figure 4B**). We monitored BNAb retention daily for a week and observed that by day 8 post BNAb loading, over 85% of CD64 positive transduced NK cells still retained the FITC conjugated 10-1074 BNAb with strong Geometric Mean Fluorescence Intensity (**Figure 4A**). Not only could the CD16 expressing control NK cells not retain the BNAb long-term like the CD64 transduced NK cells, but all of FITC conjugated 10-1074 BNAb had fallen off the control NK cells by the next day after pre-loading (**Figure 4B**). The trend of CD16 expressing control NK cells dropping the BNAb quickly was consistent across experiments and indicated that our CD64 expressing genetically modified NK cells have a substantial advantage in long-term BNAb binding (**Figure 4C**).

**Figure 4.**
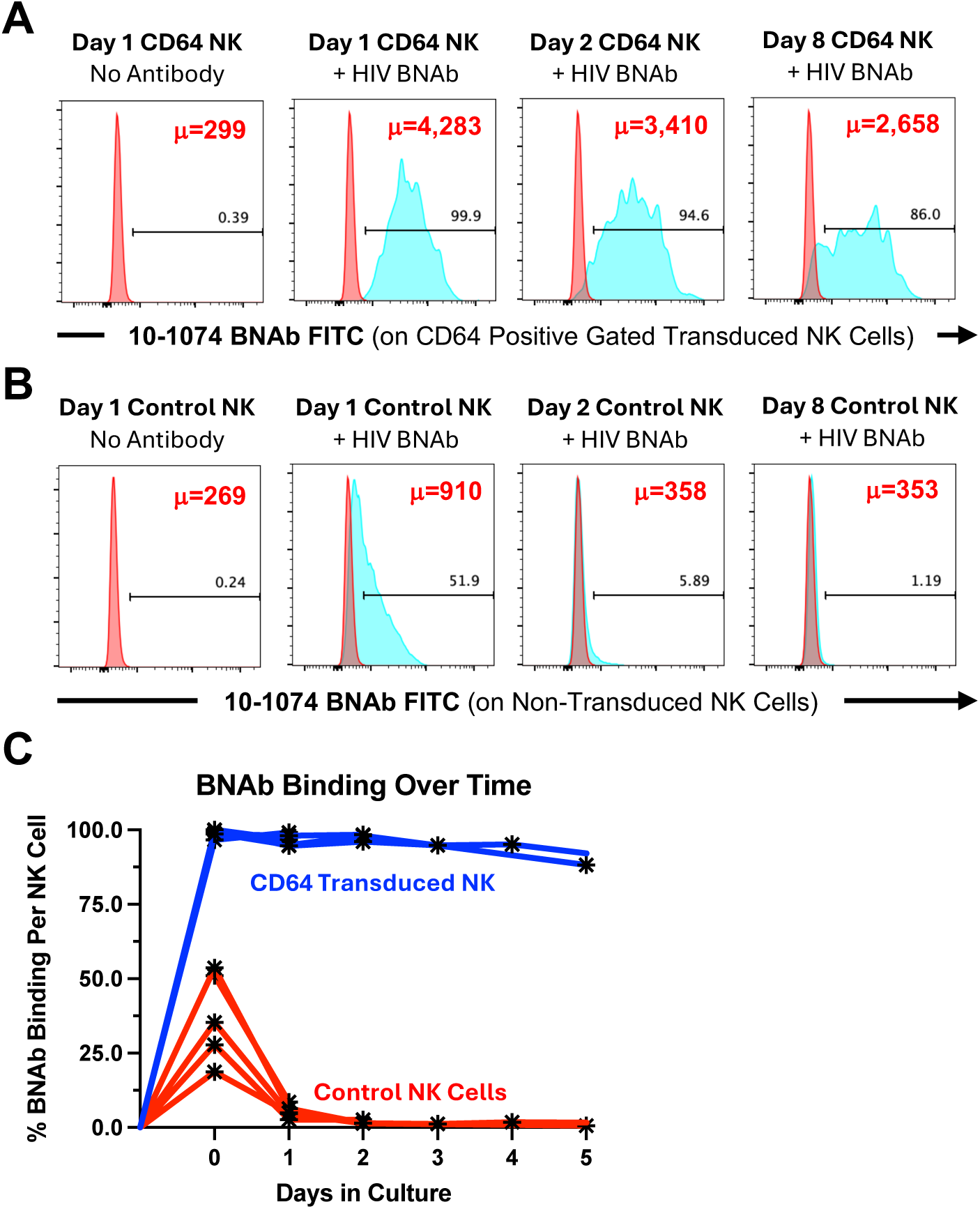
CD64 Transduced NK cells Retain HIV-specific BNAbs Over A Much Longer Time Frame Than Control NK cells in a Long-term Assay of BNAb Binding. **(A-C)** Control and CD64 Transduced NK cells were prepared as in Figure 1. On Day 7 post transduction, Control or CD64 transduced NK cells were pre-loaded with 50 μg/ml of FITC Conjugated HIV-specific 10-1074 BNAb for 15 minutes. After washing, NK cells were stained for CD64 Surface Expression and HIV-Specific BNAb fluorescence by Flow Cytometry daily over a two-week time frame to measure Long-term BNAb retention. **(A-B)** A representative experiment showing the successful retention of HIV-specific BNAbs by **(A)** CD64 Lentiviral Transduced NK cells compared to **(B)** Control NK cells over time. The Geometric Mean Fluorescence Intensity of HIV-specific 10-1074 FITC BNAb retention is shown in upper right hand part of each histogram in Red. **(C)** Composite graph showing the enhanced capacity of CD64 Lentiviral Transduced NK cells (shown in blue) to retain HIV-specific BNAbs over time on the cell surface as compared to Control NK cells (shown in red) in a Long-term binding assay (n=5).

### CD64 Transduced NK cells Can Retain HIV-specific BNAbs and Mediate Targeted ADCC Against Autologous HIV-1 Infected CD4^+^ T cells

To test if the long-term retention of HIV-specific BNAbs by CD64 genetically modified NK cells would enhance their capacity to seek out and destroy HIV-1 infected target cells through ADCC, we utilized our previously described autologous assay system for measuring both NK mediated direct cytotoxicity [37, 39] as well as ADCC [40, 41]. As target cells, we isolated CD4^+^ T cells from peripheral blood of the same individuals we isolated NK cells from and infected them with the IIIB CXCR-4 tropic HIV-1 lab adapted HIV-1 isolate. We have previously described that the IIIB isolate is strongly resistant to direct NK cytotoxicity due to having lost the ability to downregulate MHC-Class 1 proteins [37] and therefore can only be killed through ADCC. However, IIIB retains the ability to downregulate CD4 allowing for enrichment of the infected CD4^+^ T cells through a CD4 depletion column (**Figure 5A**) to achieve high levels of HIV infection as described previously [39, 47]. As effector cells, we prepared control and CD64 transduced effector NK cells from the same individual and FACS sorted the transduced NK cells to achieve a high level of CD64 surface expression for our ADCC experiments (**Figure 5B**). To test HIV-specific antibody directed targeting of HIV infected cells, we pre-loaded FACS Sorted CD64 transduced or control NK cells with a combination of the HIV-specific BNAbs 10-1074 and 3BNC117. After washing off any unbound BNAbs, we incubated pre-loaded NKs in the presence or absence of Autologous uninfected or HIV-1 infected CD4^+^ T cells for 4 hours in a standard CD107a degranulation assay as measured by flow cytometry.

**Figure 5.**
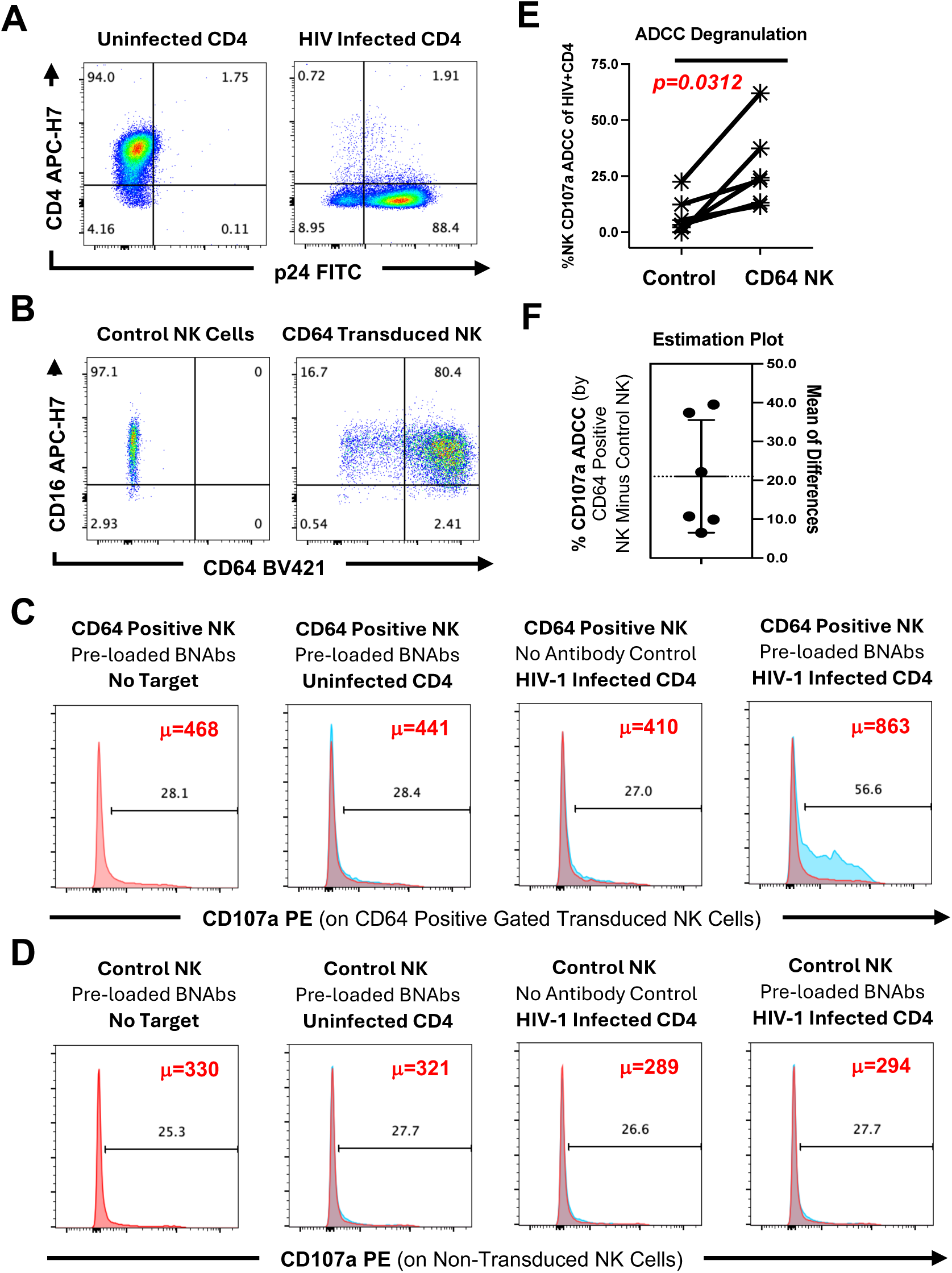
CD64 Transduced NK cells Can Retain HIV-specific BNAbs and Degranulate Against Autologous HIV-1 Infected CD4+ T cells. **(A)** CD4^+^ T cells were isolated from PHA-p stimulated PBMC and then mock or HIV-infected with 30 ng p24 containing Supernatant of IIIB CXCR-4 Tropic HIV-1 Lab Adapted HIV-1 Isolate that is resistant to direct cytotoxicity for 4 days and then Negatively Selected through a CD4 Depletion Column (Miltenyi Corporation) to Enrich p24 infected cell purity. **(B)** In parallel, Control and CD64 Transduced NK cells were prepared as in Figure 1 from the same donor PBMC and FACS sorted to achieve high CD64 surface expression (greater than 80% CD64+). On Day 7 post transduction, FACS Sorted CD64 Transduced or Control NK cells were pre-loaded with 100 μg/ml total combination of equal parts HIV-specific 3BNC117 and 10-1074 BNAbs. After washing, NK cells were incubated in the presence of Autologous Uninfected or HIV-1 infected Enriched CD4+ T cells at a 1:3 effector to target cell ratio for 4 Hours. CD107a Degranulation by **(C)** CD64+/CD56+/CD3-Transduced NK Cells or **(D)** CD56+/CD3-Control NK cells (Blue Histogram) is shown as compared to No Target Control condition (Red Histogram). The Geometric Mean Fluorescence Intensity of CD107a Degranulation is shown in upper right-hand part of each histogram in Red. **(E)** Composite graph showing statistically greater capacity of CD64 Lentiviral Transduced NK cells to retain HIV-specific BNAbs and degranulate against HIV-1 infected CD4+ T cells in a 4 Hour ADCC Degranulation Assay (n=6). **(F)** Estimation plot shows the difference in CD107a Degranulation between HIV-BNAb Pre-loaded CD64 Lentiviral Transduced NK cells and HIV-BNAb Pre-loaded Control NK cells when incubated with HIV-1 infected CD4+ T cells in a 4 Hour ADCC Degranulation Assay (n=6). Background levels of NK-mediated CD107a Degranulation against Uninfected CD4^+^ T cells was subtracted to ensure CD107a Degranulation is HIV-1 specific in Composite graphs and statistical analysis.

As shown in **Figure 5C**, CD64 positive transduced NK cells pre-loaded with the HIV-specific BNAbs were able to mediate strong ADCC CD107a degranulation against HIV-1 infected, but not uninfected, autologous CD4^+^ T cells. By contrast, control NK cells were not able to retain the HIV-specific BNAbs long enough to mediate ADCC degranulation against HIV-1 infected CD4^+^ T cells above background (**Figure 5D**). In the absence of antibody, CD64 positive Transduced NK cells were incapable of degranulating against HIV-1 infected autologous CD4^+^ T cells through direct cytotoxicity (**Figure 5C**), presumably due to the inability of the IIIB viral isolate to downregulate MHC-1 proteins as shown previously [37]. Importantly, the ability of BNAb pre-loaded CD64 positive NK cells to mediate ADCC degranulation was HIV-specific as no CD107a degranulation was observed against uninfected CD4^+^ T cells, and was also significant (*p=0.0312*, n=6) when compared to control NK cells (**Figure 5E and F**).

We next investigated if CD64 transduced NK cells could hold onto HIV-1 specific BNAbs for longer stretches of time and mediate cytotoxicity of HIV-1 infected autologous CD4^+^ T cells in a 24 hour HIV-1 elimination assay. We prepared control and CD64 transduced effector NK cells and pre-loaded them with HIV-1 specific BNAbs as described above. As targets, we isolated activated autologous CD4^+^ T cells but this time infected them with the NL4-3 CXCR-4 Tropic HIV-1 lab adapted HIV-1 Isolate rather than IIIB. NL4-3 retains the ability to downregulate MHC-Class 1 proteins [37] and is therefore susceptible to both to direct NK cytotoxicity and ADCC allowing us to compare these two killing pathways simultaneously. We then incubated the uninfected or NL4-3 infected CD4^+^ T cells for 24 hours in the presence or absence autologous CD64 transduced or control NK cells (without antibody or after pre-loading with HIV-specific BNAbs) and measured HIV elimination by measuring the frequency of intra-cellular p24 positive cells by flow cytometry staining for the HIV-1 p24 capsid protein.

Using a high MOI of infection where almost 50% of CD4^+^ T cells were HIV-1 infected (p24 positive), we observed that both control and CD64 transduced NK cells were able to mediate direct cytotoxicity of NL4-3 infected CD4^+^ T cells in the absence of antibody (41% versus 37% reduction in p24 positive events, respectively) (**Figure 6A**). However, only CD64 positive transduced NK cells were able to retain the HIV-specific BNAbs during the course of the 24 hour assay and mediate strong ADCC elimination of HIV-1 infected autologous CD4^+^ T cells above direct cytotoxicity (61% total reduction in p24 positive events when ADCC is combined with direct cytotoxicity compared to 37% with direct cytotoxicity alone) (**Figure 6A**). In contrast, control NK cells were not able to retain the HIV-specific BNAbs long enough to mediate any ADCC elimination above background against HIV-1 infected CD4^+^ T cells (42% total reduction in p24 positive events when ADCC is combined with direct cytotoxicity compared to 41% with direct cytotoxicity alone) (**Figure 6A**). The ability of CD64 positive transduced NK cells to mediate elimination of HIV-1 infected CD4^+^ T cells through direct cytotoxicity or ADCC was completely HIV specific as no reduction was observed on uninfected CD4^+^ T cells in any condition (**Figure 6A**). The enhanced ADCC elimination of HIV-1 infected CD4^+^ T cells mediated by CD64 Transduced NK cells when pre-loaded with HIV-specific BNAbs was reproducible (**Figure 6B**), and also observed when a low MOI of HIV was used (1-5% of CD4^+^ T cells were HIV-1 p24 positive infected, data not shown). While Control NK cells could not retain the HIV-specific BNAbs long enough to trigger ADCC elimination of HIV-1 infected CD4^+^ T cells, they were able to mediate direct cytotoxicity at equivalent levels as CD64 Transduced NK cells (**Figure 6C**).

**Figure 6.**
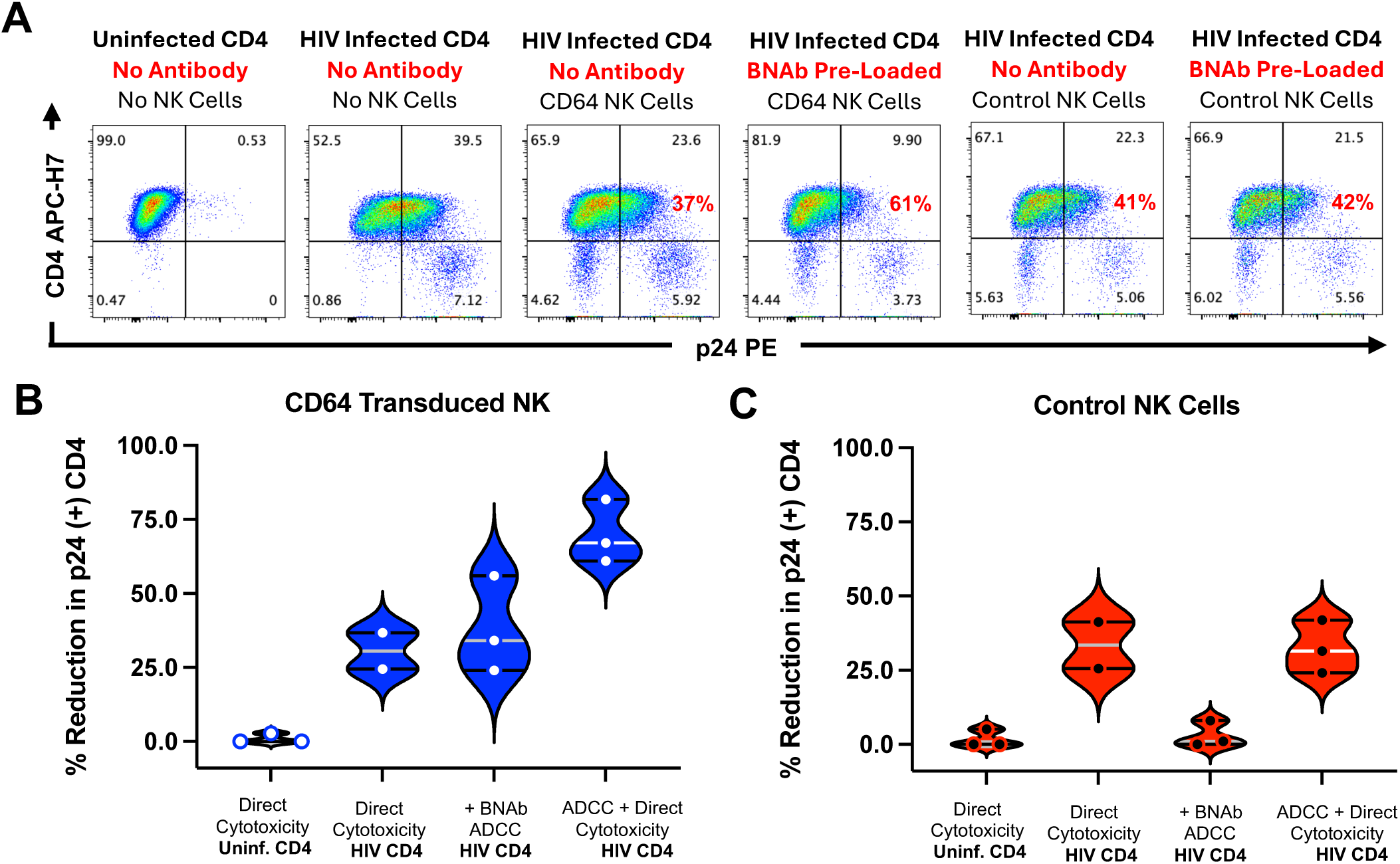
CD64 Transduced NK cells Can Retain HIV-specific BNAbs and Eliminate p24 Positive Autologous HIV-1 Infected, but not Uninfected, CD4+ T cells Through ADCC. **(A-C)** CD4^+^ T cells were isolated from PHA-p stimulated PBMC and then mock or HIV-infected for 4 days with 30 ng p24 containing Supernatant of NL4-3 CXCR-4 Tropic HIV-1 Lab Adapted HIV-1 Isolate that is susceptible to both direct cytotoxicity and ADCC. In parallel, Control and CD64 Transduced NK cells were prepared from the same individual PBMC as in Figure 1. **(A)** On Day 7 post transduction, Control or CD64 transduced NK cells were incubated alone or pre-loaded with 100 μg/ml total combination of equal parts HIV-specific 3BNC117 and 10-1074 BNAbs. After washing to remove unbound BNAbs, uninfected or HIV-1 infected CD4^+^ T cells were incubated in the presence or absence of Autologous CD64 transduced or Control NK cells for 24 Hours (with and without BNAb pre-loading) to measure ADCC or Direct Cytotoxicity, respectively. Intracellular staining for HIV-1 Capsid p24 was then carried out along with CD4 surface staining for all samples. The Percentage Reduction in p24 positive events is shown in upper right-hand part of each histogram in Red. **(B-C)** Composite graph showing the enhanced capacity of **(B)** CD64 Lentiviral Transduced NK cells to retain HIV-specific BNAbs and eliminate HIV-1 infected, but not uninfected, Autologous CD4^+^ T cells through ADCC in a 24 Hour HIV Elimination Assay as compared to Control NK Cells (n=3). To calculate ADCC-mediated cytotoxicity of HIV-1 infected Autologous CD4^+^ T cells, directed cytotoxicity (in absence of antibody) was subtracted.

## Discussion

Here, we have shown that peripheral blood NK cells can be successfully transduced to express the CD64 protein to high levels on the cell surface for several weeks utilizing a lentivirus encoding the full length high-affinity CD64 Fc1a receptor gene. CD64 transduced NK cells can be successfully pre-loaded with HIV-1 specific BNAbs at levels that are substantially higher (approximately 10,000 Geometric Mean Fluorescence Intensity units per NK cell) than traditional NK cells only expressing the naturally expressed low affinity Fc receptor CD16. Unlike CD16 which can only retain antibody for hours, genetically modified NK cells expressing CD64 can retain HIV-1 specific BNAbs on the cell surface for multiple days in culture. After loading with HIV-specific BNAbs, CD64 Transduced NK cells can then seek out HIV-1 infected CD4^+^ T cells and lyse them through targeted ADCC while not killing bystander uninfected CD4^+^ T cells. In contrast, control NK cells only expressing low-affinity Fc receptor CD16 lack the capacity to hold onto the HIV-specific BNAbs before encountering an infected target cell and therefore cannot sustain long-term killing activity of HIV-1 infected CD4^+^ T cells through targeted ADCC. Importantly, CD64 transduced NK cells can also still retain the capacity to kill HIV-1 infected CD4^+^ T cells in the absence of antibody through traditional pathways of direct NK cytotoxicity.

The success of HIV-specific BNAbs such as 3BNC117 [48–50] and 10-1074 [46, 51] in delaying viral rebound during anti-retroviral therapy (ART) interruption has highlighted the promise of immuno-therapy approaches utilizing BNAbs to target the HIV reservoir. In addition to their neutralization capacity, animal models have identified the critical nature of the Fc portion of BNAbs in viral control [52]. Recent evidence showing the ability of BNAbs such as 3BNC117 and 10-1074 to trigger robust ADCC by heterologous NK cells against of a wide variety of HIV viral isolates [9] provide further evidence that BNAbs can complement the host antibody response in targeting infected cells expressing HIV envelope through Antibody-Dependent Cellular Cytotoxicity (ADCC). Here, we have confirmed that the HIV-specific BNAbs 3BNC117 and 10-1074 can also trigger ADCC degranulation (**Figure 5**) and ADCC lysis (**Figure 6**) of autologous HIV-1 infected CD4^+^ T cells by CD64 genetically modified NK cells.

Nevertheless, the ability of HIV-1 specific BNAbs to contribute to viral eradication through Fc mediated viral clearance will always be hampered by the negative regulatory actions of ADAM-17 that degrades CD16 from the NK cell surface of traditional NK cells after degranulation and prevents repeated ADCC activity over time. Fortunately, the high-affinity Fc Receptor CD64 lacks the proteolytic cleavage site utilized by ADAM-17 to degrade CD16, thereby preserving the long-term ADCC capacity of engineered NK cells when they express CD64. Our results showing the enhanced capacity of CD64 transduced NK cells to eliminate HIV-1 infected target cells over a 24 hour period when pre-loaded with HIV-specific BNAbs in an autologous HIV elimination assay suggests that NK cells genetically modified with CD64 can indeed retain long-term ADCC capacity (**Figure 6**) in addition to their superior capacity hold onto antibodies and use them to trigger ADCC.

Once NK cells mediate cytotoxicity against a target cell, they retain potential to mediate degranulation against additional targets [53] and differentiate into mature NK cells bearing the CD57 maturation marker [43, 45]. In contrast to T cells where CD57 is expressed upon senescence, the CD57 maturation marker is expressed on a subset of NK cells (typically 40-80% of CD56^dim^ NK cells) that correlates with age and the diversity of the NK repertoire [43, 45]. NK cells expressing CD57 have been shown to possess higher NK cytokine production, [54] and enhanced CD16 signaling [45]. Here, we have identified that not only could we readily transduce CD57 positive mature NK cells utilizing a lentivirus encoding the full length high-affinity CD64 Fc1a receptor gene, but we could maintain the CD57 maturation status of our isolated peripheral blood NK cells over time in culture (**Figure 2**). This is important for the potential use of CD64 genetically modified NK cells in future HIV eradication strategies as we have recently shown that the CD57 positive NK fraction represents the pre-dominant NK effector population in the peripheral blood that can mediate ADCC against HIV-1 infected autologous CD4^+^ T cells in both Control Donors and PLWH [40].

In addition to HIV CURE directed research, CD64 genetically modified NK cells may have therapeutic potential against a wide variety of tumors or other pathogens, simply by exchanging the BNAb pre-loaded onto their cell surface. Nevertheless, the relevance of this work to HIV Eradication strategies in particular is profound. Firstly, CD64 genetically modified NK cells pre-loaded with HIV-1 specific BNAbs can eliminate HIV infected cells expressing envelope proteins through ADCC. This is critical for HIV curative strategies as HIV infected cells expressing envelope proteins represent the portion of infected cell fraction with the greatest potential to reseed the reservoir during productive infection or after the use of latent reversing agents [1–5]. In addition, CD64 genetically modified NK cells retain their capacity to eliminate infected cells through direct cytotoxicity which allows them to also target HIV-1 infected cells lacking envelope but exhibiting patterns of stress after infection. We have previously shown that NK lysis of HIV-1-infected autologous CD4^+^ primary T cells through the direct cytotoxicity pathway by interferon-alpha-activated NK cells requires the NKp46 and NKG2D activating receptors [39]. As our CD64 genetically modified NK cells express NKp46 on the cell surface (**Supplementary** Figure 1), future experiments can confirm if NKp46 is also involved in direct recognition of HIV infected cells by our CD64 genetically modified NK cells.

In addition to targeted ADCC, CD64 genetically modified NK cells can be combined with other NK immuno-therapy strategies such as BiKEs and TriKES [55–57] or Lymph node homing receptor strategies [58, 59] to direct these CD64 genetically modified NK cells to HIV reservoir sites where they can carry the broadly neutralizing antibodies needed to kill HIV infected cells. Taken together, these antibody-loaded CD64 expressing chimeric NK cell strategy or “NuKES” (<u>NK</u> <u>E</u>nhancement <u>S</u>trategy) are attractive for future clinical development as a potential autologous CAR primary NK eradication strategy against HIV due to their capacity to retain HIV-specific BNAbs, resist negative regulation, and still mediate HIV-targeted ADCC for prolonged periods after antibody-loading.

## Materials and Methods

### Subject Criteria and Clinical Assessment

16 Uninfected control donors and 8 ART-Suppressed People Living with HIV (PLWH) were enrolled from the greater Philadelphia metropolitan area according to Informed Consent Principles. All participants were recruited according to IRB guidelines approved by the Institutional review boards of the University of Pennsylvania, Presbyterian Hospital, Philadelphia FIGHT, and The Wistar Institute (IRB Approval Number 2110176). All ART-suppressed PLWH received anti-retroviral therapy with undetectable viral replication (below 50 copies viral RNA/mL) for a period of at least one year prior to recruitment and were recruited with CD4^+^ T cell counts above 350 cells/ microliter at the time of draw.

### Multi-color Flow Cytometry

To assure viability, NK cells were stained with Aqua Live/Dead (ThermoFisher Scientific, Waltham, MA) in 1X Phosphate Buffered Saline (PBS) for 30 minutes at 4°C in the dark followed by surface staining. Cells were stained with antibodies to phenotypic and functional markers in PBSA Buffer (1X PBS plus 0.09% Sodium Azide) with 25% volume of Brilliant Staining Buffer (BD Biosciences, San Jose, CA) for 15 minutes at room temperature in the dark, washed twice and then fixed for 5 minutes at 4°C with CytoFix/CytoPerm Buffer (BD Biosciences). The following surface antibodies were used at the manufacturer recommended dilution of 0.25 μg antibody per million cells with the exception of CD57 APC which was tittered down 1/5 in order to be on scale: HLA-DR BUV395 (BD Biosciences, Clone: G46-6), NKp46 BV650 (BD Biosciences, Clone: 9E2), CD64 BV421 (BD Biosciences, Clone: 10.1), CD38 FITC (BD Biosciences, Clone: HB7), CD107a PE (BD Biosciences, Clone: H4A3), CD56 PERCP Cy5.5 or RB705 (BD Biosciences, Clone: B159), NKG2A PE-Cy7 (Beckman Coulter, Pasadena, CA. Clone: Z199), CD57 APC (BioLegend, San Diego, CA. Clone: QA17A04), CD16 APC-H7 (BD Biosciences, Clone: 3G8), CD3 BUV809 (BD Biosciences, Clone: UCHT1), CD4 APC-H7 (BD Biosciences, Clone: RPA-T4). Intra-cellular staining for p24 PE (Beckman Coulter, Clone: Kc57) was carried out in 1X Perm/Wash Buffer (BD Biosciences) for 15 minutes at room temperature in the dark. For HIV-1 envelope surface staining of gp120, HIV-1 infected CD4^+^ primary T cells were incubated with 50 μg/mL of the FITC Conjugated 10-1074 BNAb during surface staining. A minimum of two hundred thousand events were collected on a BD FACSymphony A5 SE Flow Cytometer and samples analyzed using FlowJo software (version 10.10, Tree Star, Ashland, OR).

### CD64 Transduction Experiments

CD64 Lentivirus was prepared from 293T cell transfection and ultra-centrifuged as previously described [60]. Briefly, 293T cells were seeded in a T175 flask containing Dulbecco’s Modified Eagle’s Medium (DMEM) supplemented with 10% FBS and 1% penicillin/streptomycin, and then incubated at 37 °C with 5% CO2. After two passages, 22 million cells were seeded in a T175 flask containing Opti-MEM I Reduced Serum Medium supplemented with 5% FBS, 0.5% penicillin/streptomycin, GlutaMAX supplement, and Sodium Pyruvate (Packaging Medium). Following a 24-hour incubation period, when the cell confluency reached 95%, the cells were transfected with Lipofectamine 3000 along with 21 μg of Gag/pol plasmid, 21 μg of PRSV/Rev plasmid, 3.5 μg of Cocal-g plasmid, and 31.5 μg of CD64 plasmid, all dissolved in Opti-MEM medium. Six hours post-transfection, the media was replaced with warm packaging media. The supernatant was collected 24- and 48 hours post-transfection and carefully centrifuged at 2000 rpm for 10 minutes at room temperature to remove cellular debris. Subsequently, the supernatant was centrifuged in a Beckman Coulter Optima XE-90 ultracentrifuge with a Type 70 Ti fixed-angle rotor at 100,000 g for 2 hours at 4 °C to concentrate the virus 100X. Finally, the concentrated virus was aliquoted and stored at -80 °C for long-term storage. NK cells were purified by negative magnetic bead separation (Miltenyi Corporation) to 99% purity from the peripheral blood of Control Donors (Whole Blood Draw) or Persons Living with HIV (Apheresis Preparation) and immediately activated with 100 u/mL of IL-2, 100 ng/mL of IL-15 and 100 ng/mL of IL-21 for two days. 1x10^6^ Activated NK cells were then either mock or virus transduced with 100 μl of 100X concentrated CD64 expressing Lentiviral particles via Spinfection for two hours at 1800 rpm. Media was changed on NK cells with fresh cytokines added every few days when NK cells were stained for Flow Cytometry to assess CD64 surface expression.

### BNAb Binding and Retention Experiments

Control and CD64 Transduced NK cells were prepared as described above. On Day 7 post transduction, Control or CD64 transduced NK cells were pre-loaded with 50 μg/ml of FITC Conjugated HIV-specific 10-1074 BNAb for 15 minutes at room temperature in the dark and then washed three times to remove any unbound BNAb. NK cells were stained for CD64 Surface Expression and FITC Conjugated 10-1074 BNAb fluorescence by Flow Cytometry before and after washing to measure short-term BNAb Binding. To measure Long-term HIV-specific BNAb retention, Control or CD64 transduced NK cells were pre-loaded with 50 μg/ml of FITC Conjugated 10-1074 BNAb for 15 minutes at room temperature in the dark, washed three times and then stained daily for two weeks by Flow Cytometry for CD64 Surface Expression and FITC Conjugated 10-1074 BNAb fluorescence.

### HIV-1 Infection

To trigger activation of CD4^+^ primary T cells, PBMC were incubated in the presence of 10 μg/ml PHA-p (Sigma Aldrich Corporation, St. Louis, MO) and 100 IU/ml hIL-2 (PeproTech, Rocky Hill, NJ) for 72 hours. CD4^+^ primary T cells were then isolated to 99% purity by positive selection using CD4 magnetic bead isolation kit as described by the manufacturer (Miltenyi Corporation). Activated CD4^+^ T cells at 1x10^6^/mL were spinfected at 1800 rpm for two hours with 30 ng of p24 containing supernatant of the CXCR4-tropic HIV-1 isolate IIIB or NL4-3 as previously described [37]. HIV-1 infected cells were enriched to greater than 75% p24 positivity utilizing CD4 depletion column (Miltenyi Corporation) to remove uninfected cells as previously described [39]. HIV-1 infection was determined four days later by measuring intra-cellular levels of the p24 capsid protein and surface levels of gp120 envelope levels by Flow Cytometry using the 10-1074 FITC BNAb. All viral strains were generated and tittered by the University of Pennsylvania Centers for AIDS Research (CFAR).

### NK CD107a Degranulation Assay Against Autologous HIV-1 infected CD4+ T cells

Control and CD64 Transduced NK cells were prepared as in Figure 1. On Day 7 post transduction, Control or CD64 transduced NK cells were FACS sorted to achieve uniform CD16 expression (over 96%) and high CD64 surface expression on Transduced NK cells (greater than 80% CD64 positive). In parallel, Uninfected and HIV-1 enriched autologous CD4+ T cells were prepared as above from the same donor PBMC. FACS sorted Control or CD64 transduced NK cells were then pre-loaded with 100 μg/ml total combination of equal parts HIV-specific 3BNC117 and 10-1074 BNAb for 15 minutes. After washing to remove any unbound BNAb, 1x10^6^ NK cells were incubated in the presence or absence (No Target Control) of Autologous Uninfected or HIV-1 infected Enriched CD4^+^ T cells at a 1:3 effector/target ratio along with 20 μl anti-CD107a monoclonal antibody in a 200 μl total volume for four hours. Following the degranulation assay, samples were stained by Flow Cytometry with antibodies to NK cell phenotypic markers as described above. CD107a Degranulation was determined on CD64+/CD56+/CD3-gated Transduced NK Cells or **(D)** CD56+/CD3-Control NK cells (Blue Histogram) and overlayed over No Target Control condition (Red Histogram). The Geometric Mean Fluorescence Intensity of CD107a Degranulation is shown in upper right-hand part of each histogram in Red. Background levels of NK-mediated CD107a Degranulation against Uninfected CD4^+^ T cells was subtracted to ensure CD107a Degranulation is HIV-1 specific in Composite graphs and statistical analysis.

### NK HIV-Elimination Assay Against Autologous HIV-1 infected CD4+ T cells

Control and CD64 Transduced NK cells were prepared as in Figure 1. On Day 7 post transduction, Control or CD64 transduced NK cells were then pre-loaded with 100 μg/ml total combination of equal parts HIV-specific 3BNC117 and 10-1074 BNAbs for 15 minutes followed by washing to remove any unbound BNAb. In parallel, Uninfected and HIV-1 enriched autologous CD4^+^ T cells were prepared as above from the same donor PBMC. 1x10^6^ Uninfected and HIV-1 enriched autologous CD4^+^ T cells were incubated in the presence or absence of BNAb pre-loaded Control or CD64 Transduced NK cells at a 1:5 effector/target ratio along in a 200 μl total volume for 24 hours. Following the degranulation assay, live/dead determination with carried out using Aqua Live/Dead kit, then samples were stained by Flow Cytometry with antibodies to T cell and NK cell phenotypic markers, followed by intra-cellular staining for the HIV-1 p24 Capsid protein. The presence HIV-1 infected p24 positive CD4^+^ T cells was determined on CD3+/CD56-gated T cells and compared to No Effector Control condition. The percentage reduction of HIV-1 p24 positive CD4^+^ T cells is shown in upper right hand quadrant of each dot plot in red and was calculated by dividing the number of HIV-1 p24 positive CD4^+^ T cells remaining after overnight incubation with autologous Control or CD64 Transduced NK cells by the total percentage of HIV-1 p24 positive CD4^+^ T cells present after 24 hours in the absence of NK cells (followed by then subtraction from 1). To calculate ADCC specific lysis, the elimination of HIV-1 p24 positive CD4^+^ T cells by Control or CD64 Transduced NK cells in the absence of antibody (Direct Cytotoxicity) was subtracted from total lysis when Control or CD64 Transduced NK cells were pre-loaded with HIV-1 specific BNAbs.

### Statistical Analysis

All graphic presentations were performed with Prism 10 software (GraphPad Software, La Jolla, CA) and displayed as median with interquartile range. Statistical analyses of two groups was performed using a paired, non-parametric Wilcoxon Signed-Rank Test with a two-tailed p-value for all experiments with over 6 data points or more. No statistical analysis was attempted on data with sample sizes of 5 experiments or less. No data was purposefully excluded from the analysis and all missing data from any subjects is due to technical issues with the assay. Due to limited sample size, reported p-values are not adjusted for multiple testing.

## Data Availability Statement

The data that support the findings of this study are available from the corresponding author upon reasonable request.

## Ethics Approval Statement

All participants were recruited according to IRB guidelines approved by the Institutional review boards of the University of Pennsylvania, Presbyterian Hospital, Philadelphia FIGHT, and The Wistar Institute (IRB Approval Number 2110176). All authors agree with this submission and this article is not currently submitted elsewhere.

## Patient Consent Statement

Uninfected control donors and ART-Suppressed Persons Living with HIV (PLWH) were enrolled from the greater Philadelphia metropolitan area according to Informed Consent Principles.

## Permission to Reproduce Materials from Other Sources

No materials of any kind were used from outside sources including unpublished references.

## Clinical Trial Registration

Not Applicable.

## Acknowledgments

We would like to acknowledge Sabine Baxter at the University of Pennsylvania Cell Center for her contribution in conjugating the FITC fluorochrome to the HIV-specific 10-1074 BNAb.

## Author Contributions

CT (Performed CD64 Lentiviral Transduction experiments, BNAb loading experiments, and functional ADCC CD107a and HIV Elimination Assays, performed statistical analysis, co-wrote the manuscript), AOO, LDL, and HK (Produced CD64 Lentiviral Preps including 293T cell Lentiviral transfection, harvesting and ultra-centrifugation), JLR (Provided CD64 Lentivirus and transduction expertise in initial phases of the project, helped conceptualize the study idea, edited the manuscript), LJM (Designed and coordinated the study, provided statistical support, co-wrote the manuscript).

---

**Supplementary Figure 1.**
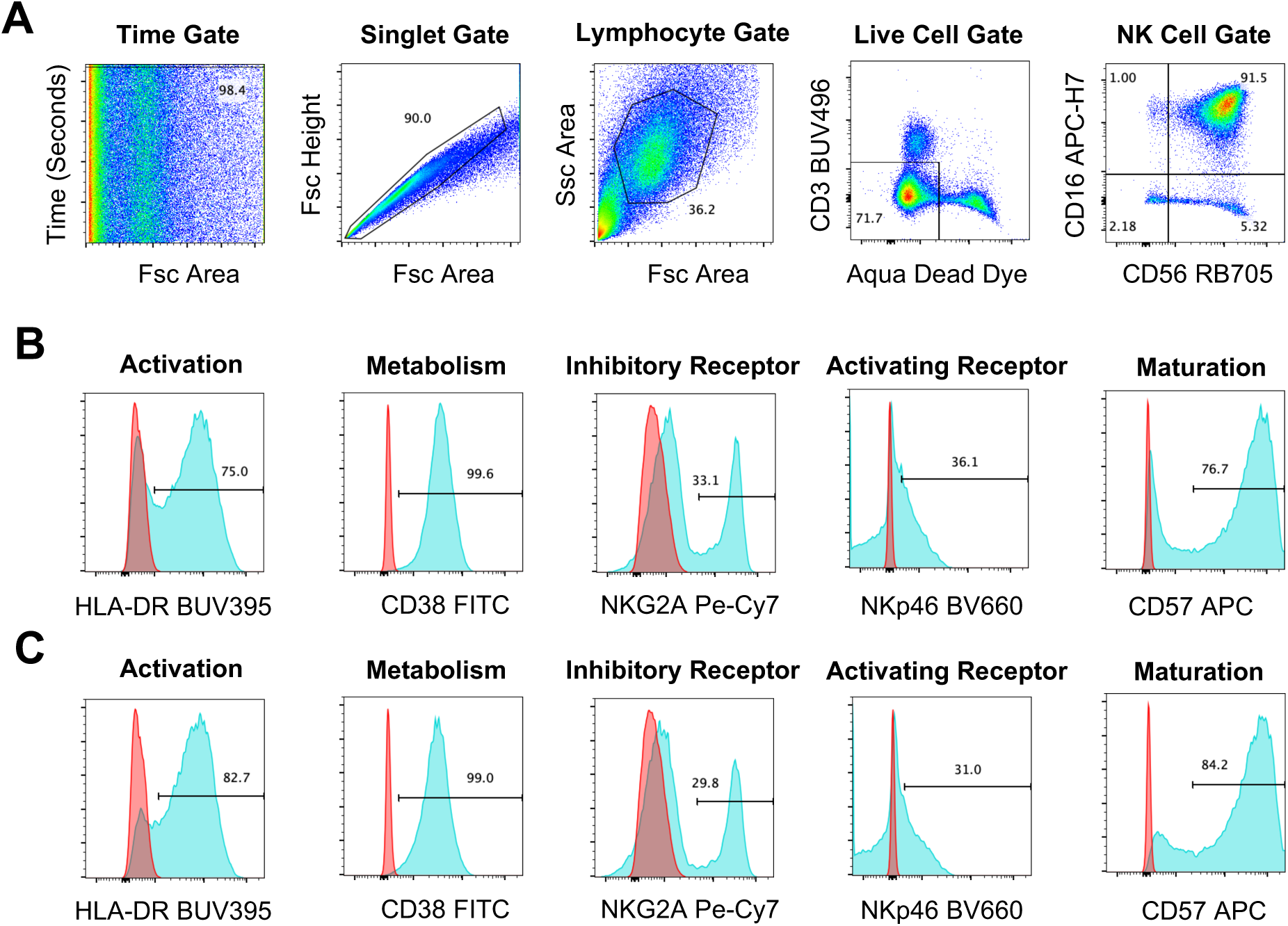
Gating Strategy and Phenotypic Analysis of CD64 Transduced NK cells Isolated from the Peripheral Blood. **(A)** CD64 Transduced NK cells were prepared as in Figure 1. On Day 7 post transduction, CD64 transduced NK cells were stained for Phenotypic Expression of surface markers by Flow Cytometry. At least 100,000 cells per condition were acquired by Flow Cytometry on FACSymphony A5 SE. NK cells were analyzed with FlowJo 10.10.0 software and gated according to Time Gate (to remove bubbles), Singlet Gate (to remove doublets), a Lymphocyte Gate (to remove Myeloid cells), Live Cell Gate (to remove dead cells and CD3 T cells), with staining for CD56 and CD16 to identify NK^dim^ Cytolytic cells. **(B-C)** Typical surface phenotype (Blue Histogram) of **(B)** Control NK cells or **(C)** CD64 Transduced NK cells compared to FMO Stain (Red Histogram) one week post spinfection with CD64 encoding Lentivirus.

