## Supplemental Figure 1 for "Gene-modified NK Cells Expressing CD64 and Pre-loaded with HIV-specific BNAbs Target Autologous HIV-1 Infected CD4^+^ T Cells by ADCC"

### Slide 1
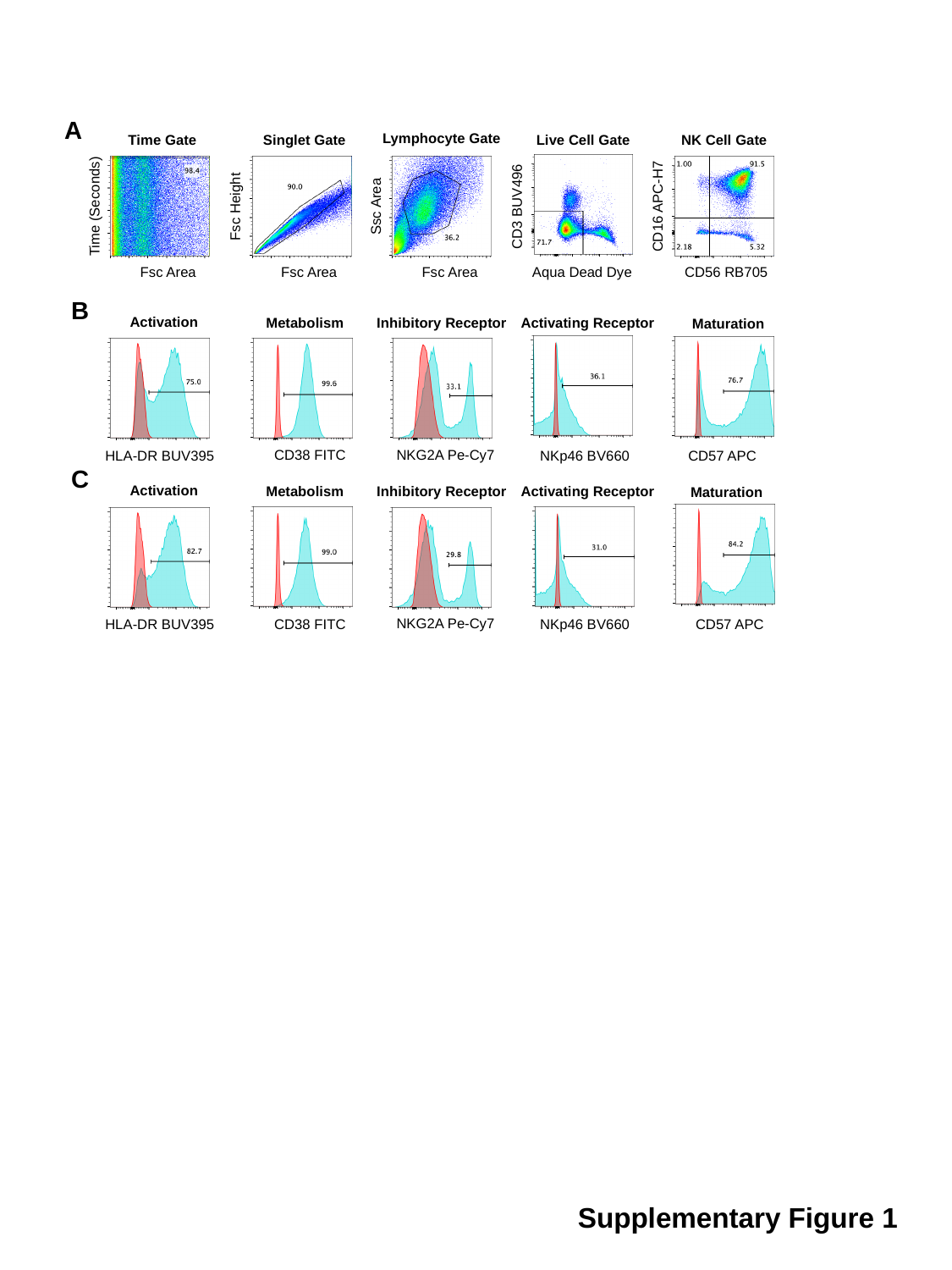

A
Lymphocyte Gate
Time Gate
Singlet Gate
Live Cell Gate
NK Cell Gate
Time (Seconds)
Fsc Height
Ssc Area
CD3 BUV496
CD16 APC-H7
Fsc Area
Fsc Area
Fsc Area
Aqua Dead Dye
CD56 RB705
B
Activation
Inhibitory Receptor
Metabolism
Activating Receptor
Maturation
NKG2A Pe-Cy7
CD38 FITC
NKp46 BV660
HLA-DR BUV395
CD57 APC
C
Activation
Inhibitory Receptor
Metabolism
Activating Receptor
Maturation
NKG2A Pe-Cy7
CD38 FITC
NKp46 BV660
HLA-DR BUV395
CD57 APC
Supplementary Figure 1
